# A novel CEST-based approach for reliably assessing skeletal muscle oxidative phosphorylation: OXCEST

**DOI:** 10.1101/2025.04.25.650537

**Authors:** Ritambhar Burman, Yuxi Pang, Asim Bag, Richard Finkel, Kirsten Ness, Puneet Bagga

## Abstract

**Purpose:** To develop and validate a novel chemical exchange saturation transfer (CEST) MRI method to map skeletal muscle OXPHOS (Oxidative Phosphorylation CEST or OXCEST).

**Theory and Methods:** Our proposed OXCEST method acquires creatine (Cr)-weighted CEST maps by applying RF saturation (B_1_) at only two frequency offsets: +1.8 ppm (targeting the Cr amine resonance) and −1.8 ppm (to calculate MTR_asym_ at 1.8 ppm). The pre-exercise MTR_asym_ is modeled as a second-order polynomial function (f) of B_0_. Next, the post-exercise alteration in MTR_asym_ is hypothesized to be affected by both an exercise-induced increase in Cr and changes in B_0_ inhomogeneity. By inputting post-exercise B_0_ values into f, the change in MTR_asym_ due to B_0_ variation alone was estimated. Thus, the Cr-related post-exercise MTR_asym_ could be isolated and quantified. OXCEST and 31P-MRS were performed in seven subjects across two sessions to compare the OXCEST-derived Cr recovery time constant (T_Cr_) with the ground-truth phosphocreatine recovery time constant (T_PCr_).

**Results:** A second-order polynomial function f could reliably describe the relationship between pre-exercise MTR_asym_ and B_0_ (R^2^=0.87±0.07 in the lateral gastrocnemius (LG); R^2^=0.98±0.01 in the medial gastrocnemius (MG); R^2^=0.96±0.03 in the soleus). The mean pre-exercise MTR_asym_ was approximately 6-7% for all muscle groups. Following exercise, MTR_asym_ increased by 11.4±4.5% in LG and 8±2.4% in MG, and showed mono-exponential recovery (R^2^>0.97). The combined T_Cr_ of LG and MG was found to be significantly correlated with T_PCr_ (R²=0.83, p=0.005).

**Conclusion:** OXCEST enables reliable assessment of post-exercise Cr recovery and demonstrated strong agreement with 31P-MRS.

## 1 INTRODUCTION

Mitochondrial oxidative phosphorylation (OXPHOS) is a critical metabolic pathway that produces adenosine triphosphate (ATP), the primary energy currency. OXPHOS capacity varies significantly across populations, with endurance athletes exhibiting considerably greater mitochondrial activity compared to sedentary lean or overweight individuals.^1^ Conversely, compromised mitochondrial OXPHOS is associated with various pathological conditions including neuromuscular disorders, mitochondrial diseases, and aging.^2–5^ Currently, invasive muscle biopsy represents the gold-standard method to assess OXPHOS capacity.^6^ However, biopsies have inherent limitations, including patient discomfort, risk of complications, and limited feasibility for repeated assessments over time. 31-Phosphorus magnetic resonance spectroscopy (31P-MRS) has traditionally been regarded as the MR-based gold standard technique to non-invasively quantify mitochondrial function in skeletal muscle.^7–11^ Phosphocreatine (PCr) is hydrolyzed during muscle contraction to regenerate ATP from ADP, and subsequently resynthesized during recovery phase through the creatine kinase (CK) reaction, which biochemically couples PCr and creatine (Cr).^12,13^ 31P-MRS has been increasingly used to investigate mitochondrial dysfunction in aging and mitochondrial disorders.^8,14–16^ However, the broader application of 31P-MRS is challenged by its relatively low intrinsic sensitivity, the need for specialized hardware, and lower spatial resolution compared to proton-based imaging techniques.

To address this, the foundation for proton imaging of Cr was established by Haris et al.,^17^ who identified sharp amine proton resonances of Cr at +1.8 ppm. Kogan et al.^18,19^ subsequently demonstrated that chemical exchange saturation transfer (CEST) imaging could map Cr in skeletal muscle following plantar flexion exercise with high spatial resolution. This approach was extended to in-vivo mapping of free Cr in myocardial disorders.^20^ B_0_ field inhomogeneity can bias CEST contrast measurements.^21^ To obtain B_0_-corrected MTR_asym_ (the difference between magnetization transfer ratio values at the positive offsets and the corresponding negative offsets with respect to water), a partial or full z-spectrum is often acquired.^17,22^ In the context of post-exercise recovery, computing MTR_asym_ at a single time point using multiple frequency offsets is inadvertently influenced by a time-varying CEST signal from multiple time-points as Cr concentration declines exponentially immediately after exercise. A recent report by Phillip Zhe Sun^23^ recently introduced an expedient B_0_ inhomogeneity correction algorithm for fast pH imaging that samples only two frequency offsets by exploiting the observation that MTR_asym_ can be modeled as a second-order polynomial function of B_0_ inhomogeneity.

Building on these findings, we have developed oxidative phosphorylation CEST (OXCEST) technique, which integrates the fast B_0_-correction method^23^ to the existing CEST approach to map Cr recovery following exercise in muscle, as previously described.^17–19^ Specifically, OXCEST images were acquired at only two frequency offsets: +1.8 and −1.8 ppm. Fast-B_0_ correction paradigm was then applied to evaluate pre- and post-exercise MTR_asym_. We observed that Cr recovered mono-exponentially post-exercise, consistent with the 31P-MRS-derived mono-exponential recovery kinetics of PCr.

## 2 THEORY

### 2.1 Cr mapping in skeletal muscle

A typical z-spectrum is primarily influenced by the following CEST pools: (i) direct water proton saturation (DS), the largest water signal intensity drop occurring at 0 ppm, (ii) conventional semi-solid magnetization transfer (MT) arising from the “bound” pool of water molecules cross-relaxing with protons in the free water pool, (iii) amide proton transfer (APT) at 3.5ppm, arising from the exchangeable amide protons in the backbone of mobile proteins and peptides, (iv) exchange-relayed nuclear Overhauser effect (rNOE) at −3.5ppm, from aliphatic protons of mobile macromolecules, and (v) Cr CEST at 1.8ppm, from the exchangeable amine protons of Cr.

The asymmetry analysis of the z-spectrum (defined by Equation 1, also called magnetization transfer ratio asymmetry, MTR_asym_) can separate the symmetrical contribution of DS and MT.

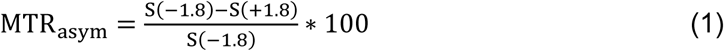

At high B_1_ (∼3 µT) used in this study, the contribution of PCr at ±1.8 ppm is negligible in skeletal muscle, and while APT and rNOE contributions at this offset are negligible in the brain at similar B_1_ levels, no such studies exist for skeletal muscle.^24–28^ Therefore, without considering B_0_ inhomogeneity effects, RF irradiation at only two frequency offsets (±1.8 ppm) is sufficient to quantify the pre-exercise CEST contribution of Cr using MTR_asym_ (MTR_asymPre_). However, as the resonance frequency in MRI is proportional to B_0_, even a minor magnetic field inhomogeneity (ΔB_0_) can cause substantial frequency shifts in z-spectrum.^21^ To correct for this B_0_ inhomogeneity, additional frequency offsets are acquired.^17,22^

Following interventions such as mild plantar flexion exercise, intramuscular pH in skeletal muscle remains relatively constant at pressures below 60% of maximal voluntary contraction (MVC) during plantar flexion,^29^ and therefore does not significantly influence the CEST contrast. Consequently, exercise-induced change in Cr level, mediated by the CK reaction, is expected to be the primary contributor to the post-exercise MTR_asym_ (MTR_asymPost_). Additional contribution to MTR_asymPost_ arises from exercise-induced motion, which perturbs B_0_,^30^ and causes post-exercise ΔB_0_ (ΔB_0Post_) to differ from the pre-exercise ΔB_0_ (ΔB_0Pre_). Given that PCr resynthesis is stoichiometrically coupled with a decrease in free Cr, a higher temporal resolution scan will enable capturing rapid changes in Cr levels. Here, we propose limiting the CEST acquisition to +1.8 and −1.8 ppm, thereby attaining a temporal resolution of 10s (5s per offset x 2 offsets).

### 2.2 Fast B_0_ correction in OXCEST

To account for B_0_ inhomogeneities, we modeled the relationship between MTR_asymPre_ and the corresponding ΔB_0Pre_ across all pixels within a muscle. A second-order polynomial function was fit using least-squares regression to obtain the three coefficients (c_0_, c_1_, c_2_):^23^

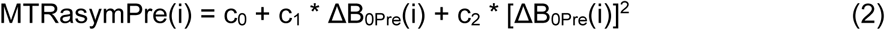

for i = 1, 2,…, N_muscle_.

Function f with fitted coefficients (c_0_, c_1_, c_2_) capture how much the MTR_asymPre_ varies as a function of ΔB_0Pre_ in a given muscle. This fitted function f is then applied to the ΔB_0Post_ values to estimate the portion of the post-exercise MTR_asym_ (MTR_asymPost_) signal that is attributable solely to exercise-induced changes in B_0_ alone (i.e. not due to actual physiological change in Cr level):

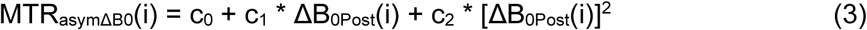

We hypothesize that the observed MTRasymPost at each pixel results from two additive effects:

_1._ A B_0_-induced change: MTR_asymΔB0_
2. A physiologic change due to change in Cr concentration: MTR_asymΔCr_ Thus, the measured MTR_asymPost_ is expressed as:

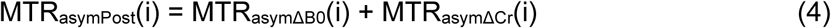

By subtracting the estimated B_0_-induced component, we isolate the signal due to Cr recovery:

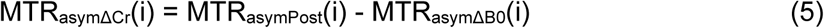

Next, for an entire muscle, MTR_asymΔCr_ can be calculated as the average of MTR_asymΔCr_ over all the pixels in the muscle.

## 3 METHODS

### 3.1 Study Design

The protocol was approved by the Institutional Review Board, with informed written consent obtained prior to the scan. Seven healthy adult participants (male: 5, mean age: 34.6 ± 6.1 years) were scanned. Each participant underwent one MR session for OXCEST and another for 31P-MRS, conducted in two separate sessions on different days on a 3T whole-body MR scanner (MAGNETOM Prisma, Siemens Healthcare, Erlangen, Germany). During each acquisition, participants performed mild plantar flexion exercise for 2 minutes at a constant rate of 45 flexions per minute inside the scanner, using an MR-compatible ergometer (Trispect, Ergospect GmbH, Innsbruck, Austria). The pedal pressure was set to 8 PSI, corresponding to 30–40% of each participant’s MVC, determined prior to scanning as the peak pressure they could tolerate. The MRI protocols for OXCEST and 31P-MRS are summarized in **Figure S1**.

### 3.2 OXCEST MRI

OXCEST data were acquired with a Tx/Rx 1/15-channel 1H knee coil (Siemens Healthcare, Erlangen, Germany). A localizer was first acquired on the right calf of each participant to center the imaging slice on the thickest section of the calf muscle. Before exercise, one image without selective saturation, a B_1_ map using a double angle method (30° and 60°),^31^ a B_0_ map using standard water-saturation shift-referencing (WASSR),^21^ and four baseline CEST images^19^ were acquired. Immediately following a 2-minute exercise, OXCEST was performed which included post-exercise CEST acquisitions (60 images at 10s temporal resolution), B_0_ map, B_1_ map, and one reference image without selective saturation. Post-exercise muscle boundaries were delineated from post-exercise CEST images. Acquisition parameters can be found in Chakraborty et al.^32^ Briefly, OXCEST imaging parameters were set as follows: TR/TE = 4.7/2.3 ms, slice thickness = 10 mm, flip angle = 10°, B_1rms_ = 3 μT, in-plane resolution = 1.25×1.25 mm^2^, matrix size = 128×128, field of view= 160×160 mm^2^. OXCEST images were acquired with saturation frequency offsets of ±1.8 ppm relative to the water resonance, achieving a temporal resolution of 10s (5s per offset × 2 offsets). Two dummy scans were included to achieve T1 steady state. For four subjects, the shimming voxel was constrained to a volume encompassing the three muscles of interest. Although MTR_asym_ is relatively stable around optimal RF power level,^33^ both pre- and post-exercise OXCEST images were corrected for minor B_1_ shifts using an inverse scaling model, assuming a linear relationship between B_1_ amplitude and CEST signal intensity.^31^

### 3.3 31P-MRS

31P-MRS was performed with a 1H/31P dual-tuned flex transmit/receive surface coil of 11cm diameter (RAPID Biomedical GmbH, Würzburg-Rimpar, Germany) using an unlocalized free induction decay (FID) sequence: measurements = 96, averages = 2, bandwidth = 5000Hz, and TR = 5s. The total data acquisition time was 15 minutes: 5 mins of baseline, 2 mins of exercise and 8 mins of post-exercise recovery. The temporal resolution of 31P-MRS was kept at 10s to match the temporal resolution of OXCEST acquisitions.

### 3.4 Data analysis

All images and data/statistical analyses were performed on MATLAB v2023b.

#### 3.4.1 OXCEST

The lateral gastrocnemius (LG), medial gastrocnemius (MG) and soleus (Sol) muscles were manually segmented on the pre-exercise reference image and the post-exercise CEST images. Since exercise can induce shifts in muscle position and subtle geometric changes, including transient hypertrophy during recovery, regions of interest (ROIs) were re-segmented on each post-exercise image until no further changes in muscle boundaries were observed across consecutive time points (usually after 2 minutes). This approach ensured accurate tracking of muscle-specific recovery dynamics. Additionally, anterior tibialis (AT) was also manually segmented. The B_0_-corrected MTR_asymΔCr_ signals, reflecting exercise-induced changes in Cr within the three muscles, were calculated as described in section 2.2. The Cr recovery time-constant (T_Cr_) for each muscle was estimated by fitting a mono-exponential decay model, given by:

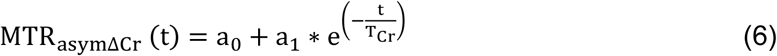

where, a_0_ is the baseline value, and a_1_ is the amplitude of exponential decay. Initial values were a_0_ = min (*MTR_asymΔcr_*), a1 = max (*MTR_asymΔcr_*) − min(*MTR_asymΔcr_*), and T*_Cr_* = 60*s*. Bound constraints were a_1_ > 0 and T_Cr_ > 0. Additionally, the pre-exercise B_0_-corrected MTR_asym_ value for each muscle was calculated from the coefficient c_0_ measured in Equation 2. The exercise-induced increase in MTRasym value was calculated from the coefficient a_1_.

To assess B_0_-correction effects, the measured MTR_asym_ (Equation 1) was averaged without B_0_ correction across all pixels in each muscle at each time point. The mean exercise-induced increase was also computed as the difference between the MTR_asymPost_ at the first time point immediately after exercise and MTR_asymPre_.

#### 3.4.2 31P-MRS

The 31P-MRS spectra were phased, baseline-corrected, and fitted with Lorentzians. The PCr peak area was quantified from each FID and normalized to the pre-exercise value. Post-exercise PCr level was expressed as a percentage of the pre-exercise level. The PCr recovery time-constant, T_PCr_, was estimated by fitting the data to a mono-exponential growth function:^34^

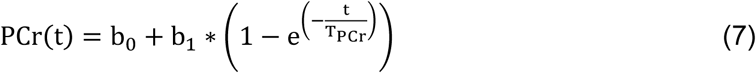

where, b_0_ is the normalized PCr level at the first post-exercise time point, and b_1_ represents the amplitude of recovery.

Additionally, the exercise-induced pH shift was estimated from the chemical shift difference between PCr and inorganic phosphate (P_i_) (δ) in 31P-MRS spectra using the adjusted Henderson-Hasselbach equation:^35^

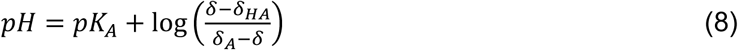

where pK_A_ = 6.75 is the dissociation constant of P_i_, δ_HA_ = 3.27 is the chemical shift of the protonated form of P_i_, and δ_A_ = 5.63 is the chemical shift of the non-protonated form of Pi.

### 3.5 Statistical Analysis

Linear regression was performed to evaluate the relationship between T_Cr_ and T_PCr_. and a two-sided p-value was calculated to test the significance of the association. Lin’s concordance correlation coefficient (CCC) was also used to assess the agreement between the two measures. Since the majority of the 31P-MRS signal acquired using a surface coil originates from the gastrocnemius muscle,^18^ the mean T_Cr_ values from the LG and MG were used for statistical comparison with T_PCr_. The coefficient of determination (R^2^) and an F-statistic was calculated from the least squares fit of a second-order polynomial relating MTR_asymPre_ to ΔB_0Pre_ in LG, MG, and Sol. 95% confidence intervals (CIs) were calculated for the predicted B_0_-corrected MTR_asymPre_ and the exercise-induced increases in MTR_asym_. Following mono-exponential fitting of Cr recovery in LG, MG and Sol, the root mean squared error (RMSE), R^2^, and standard deviation (SD) of residual noise (reported as a percentage of exercise-induced MTR_asym_ increase) were determined. A *P* < 0.05 was considered statistically significant.

## 4 RESULTS

### 4.1 Demonstration of fast B_0_ correction in OXCEST

The proposed OXCEST approach is illustrated in an axial calf muscle section of representative 35-year-old female subject in **Figure 1**. First, the LG, MG, and Sol muscles were manually segmented on a reference anatomical image of the calf (Figure 1A). Pixel-wise MTR_asymPre_ values were then calculated in the three muscles from OXCEST images (Figure 1B). The corresponding ΔB_0Pre_ map was derived from the pre-exercise WASSR scan (Figure 1C). Figure 1D displays a scatter plot of MTR_asymPre_ versus ΔB_0Pre_ in the MG muscle (orange crosses), along with the second-order polynomial fit (dotted black line) described by the following equation: MTR_asymPre_ = 5.91 + 152 x ΔB_0Pre_ – 191 x ΔB_0Pre2_ (R^2^ = 0.97, *P* < 0.001). The B_0_-corrected baseline MTR_asymPre_ value in MG was estimated by setting the corresponding ΔB_0Pre_ to 0 in the previous equation. This yielded a net value of 5.91% (95% CI: 5.74-6.08%) for MG. Similar net MTR_asymPre_ estimation was performed for LG (Figure S2A) and Sol (Figure S2B). The B_0_-corrected baseline MTR_asymPre_ values were 6.76 (95% CI: 6.37-7.15%) and 7.2% (95% CI: 7.01-7.29%) for LG and Sol, respectively. Following exercise, the muscles ROIs were re-segmented on the post-exercise CEST-weighted images to account for any movements during exercise. The segmented LG, MG, and Sol regions at the first post-exercise time point (t = 10s) are shown in Figure 1E. Pixel-wise MTR_asymPost_ values were then calculated from the OXCEST images (Figure 1F). The ΔB_0Post_ map was calculated from a WASSR scan acquired following the post-exercise OXCEST scan (Figure 1G). Figure 1H shows a scatterplot of MG MTR_asymPost_ values plotted against ΔB_0Post_ (hollow circles). The black crosses represent MTR_asymΔB0_ values predicted using Equation 3. The vertical distance between each MTR_asymPost_ and its associated MTR_asymΔB0_ represents the Cr-specific signal change (MTR_asymΔCr_) per pixel. Similar to MG, the MTR_asymPost_ was plotted against ΔB_0Post_ for LG and Sol showing exercise-related Cr increases (Figures S2A and S2B). Plots of MTR_asymPre_ against B_0Pre_ across all subjects in LG, MG, and Sol are shown in **Figures** S3 A-C. The fitted coefficients (c_0_, c_1_, c_2_) are presented in **Table S1**.

**Figure 1.**
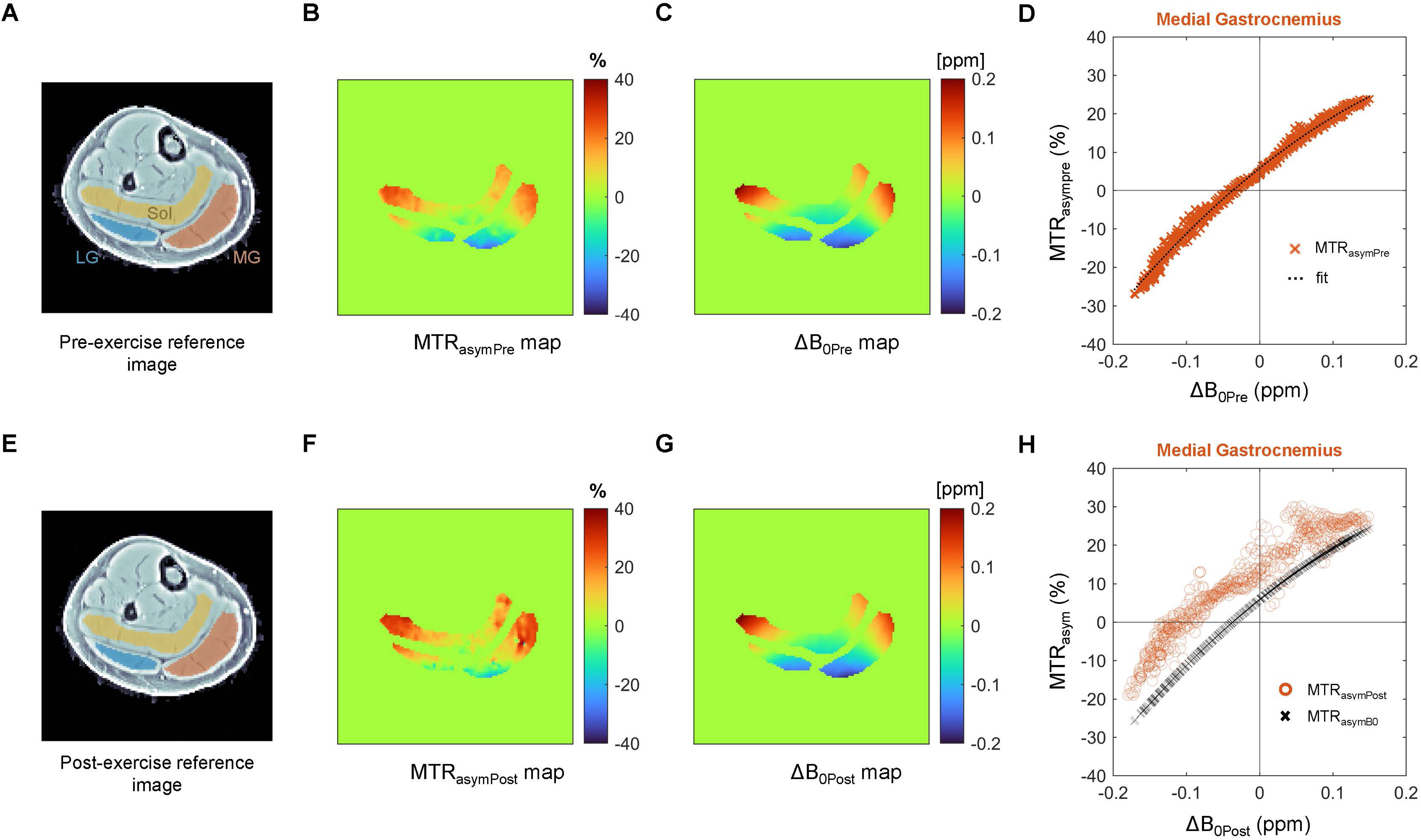
Demonstration of the Oxidative Phosphorylation Chemical Exchange Saturation Transfer (OXCEST) approach in a 35-year-old female participant. (A) Segmentation of the lateral gastrocnemius (LG), medial gastrocnemius (MG), and soleus (Sol) muscles on a pre-exercise axial anatomical calf image. (B-C) Pre-exercise magnetization transfer ratio asymmetry (MTR_asymPre_) map acquired at +1.8 ppm and the corresponding B_0_ inhomogeneity (ΔB_0Pre_) map derived from a water-saturation shift-referencing (WASSR) scan, both overlaid on the segmented muscle masks. (D) Pixel-wise regression of MTR_asymPre_ against ΔB_0Pre_ in MG; the dotted line represents the second-order polynomial fit. (E) Segmentation of the three muscles on a post-exercise axial anatomical calf image. (F-G) Post-exercise MTR_asym_ (MTR_asymPost_) and ΔB_0_ (ΔB_0Post_) maps acquired following mild plantar-flexion exercise. (H) Scatterplot of MTR_asymPost_ versus ΔB_0Post_ in MG (circles). Black crosses indicate the expected value of MTR_asym_ due solely to B_0_ changes (MTR_asymΔB0_), assuming no change in creatine (Cr) concentration. Thus, the Cr-related change in MTR_asym_ is quantified as the vertical distance between each circle and its corresponding cross, representing the ΔB_0_-corrected Cr effect.

### 4.2 Post-exercise Cr recovery mapped by OXCEST

The B_0_-uncorrected MTR_asymPost_ maps and B_0_-corrected MTR_asymΔCr_ maps for the same subject, acquired using OXCEST over a 10-minute post-exercise recovery period, are presented in **Figure 2**. At each time point, pixel intensity reflects the Cr-specific MTR_asym_ signal, with red indicating elevated values and blue indicating values approaching baseline. At 10s post-exercise, all three muscle regions display a marked increase in free Cr concentration. Over the first minute of recovery, Cr concentration rapidly decreases as it rephosphorylates to PCr via the CK reaction.

**Figure 2.**
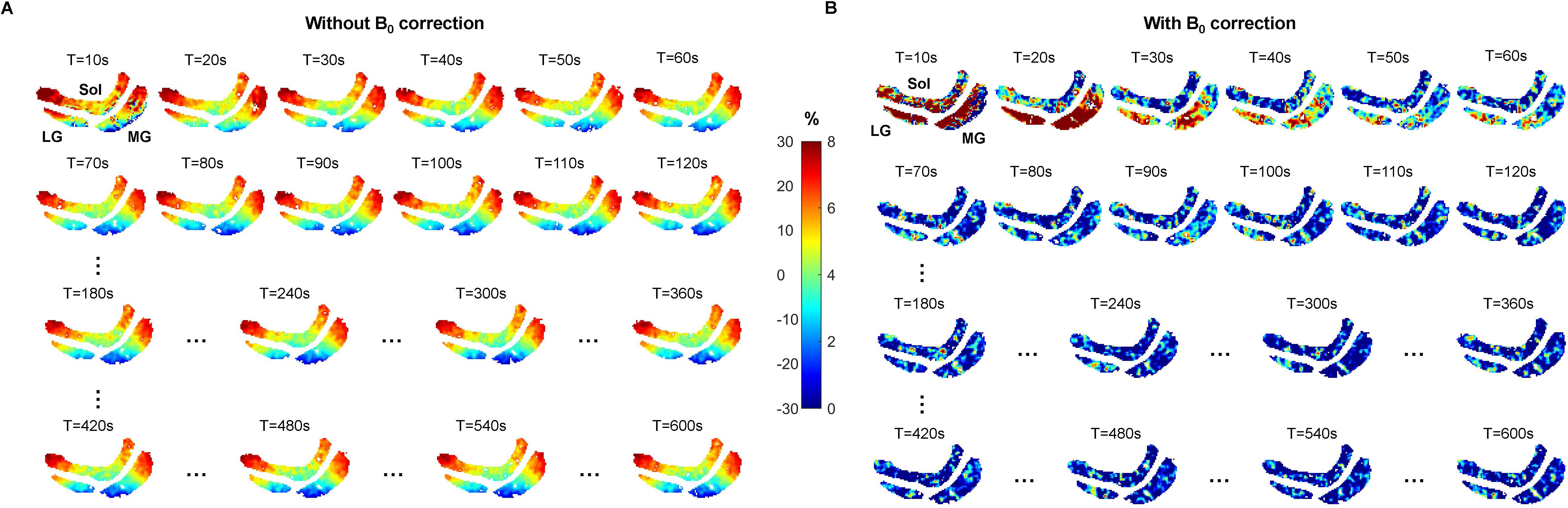
Effect of Oxidative Phosphorylation Chemical Exchange Saturation Transfer (OXCEST)-based B_0_ correction on post-exercise MTR_asym_, acquired with a temporal resolution of 10s over a 10-minute post-exercise recovery period. Each frame displays pixel-wise post-exercise magnetization transfer ratio asymmetry (MTR_asymPost_) (in %) in a representative axial slice of the lower leg encompassing the soleus (Sol), lateral gastrocnemius (LG), and medial gastrocnemius (MG) muscles: (A) without B_0_ correction, and (B) after B_0_ correction. Without correction, MTR_asym_ values often fell outside the physiologically reasonable range (e.g. negative or abnormally large values). After correction, elevated MTR_asym_ immediately after exercise (red pixels, 10-60s after exercise) reflects an increased free creatine concentration, which decays exponentially over time. This spatiotemporally resolved mapping of Cr dynamics highlights muscle-specific variations in mitochondrial OXPHOS capacity.

Table 1 contains estimated values for muscle-wise B_0_-corrrected MTR_asymPre_, exercise-induced increases in MTR_asym_, and R^2^ values of function f. The R^2^ values for the function f were found to be consistently greater than 0.8 for LG, MG, and Sol (LG: 0.87 ± 0.07, *P* < 0.001; MG: 0.98 ± 0.01, *P* < 0.001; Sol: 0.96 ± 0.03, *P* < 0.001). The B_0_-corrrected net MTR_asymPre_ values were found to be 6.03 ± 1.7% in LG, 6.76 ± 0.5% in MG, and 6.87 ± 0.4% in Sol for all the subjects. The 95% CIs for the estimated B_0_-corrrected MTR_asymPre_ were consistently narrow in MG and Sol, as well as in LG of four subjects where constrained shimming was performed. In the remaining three subjects where shimming was not constrained, the absolute B_0_ inhomogeneity exceeded 0.1 ppm in the LG only, resulting in broader 95% CIs. In the absence of registering pre- and post-exercise ROIs, the distribution of ΔB_0_ values in all three muscles are reported in **Table S2**. LG showed the largest exercise-induced increase in MTR_asym_ (11.4 ± 4.5%), followed by MG (8 ± 2.4%) and Sol (4 ± 2%), all with narrow 95% CIs. Parameters related to mono-exponential fitting of Cr recovery in LG, MG, and Sol are reported in **Table S3**. The R^2^ values of the Cr recovery mono-exponential fit were > 0.97 in both LG and MG. Net MTR_asymPre_ values in the LG, MG, and Sol muscles, as well as their post-exercise increases with and without B_0_ correction, are presented in **Table S4**. Without B_0_-correction, MTR_asymPre_ values ranged from −34% to +28.7%. Analysis of post-exercise increases in MTR_asym_ for AT muscles across all subjects, and the associated T_Cr_ are shown in **Table S5**. Both the metrics showed very high inter-subject variance for TA.

**Table 1:**
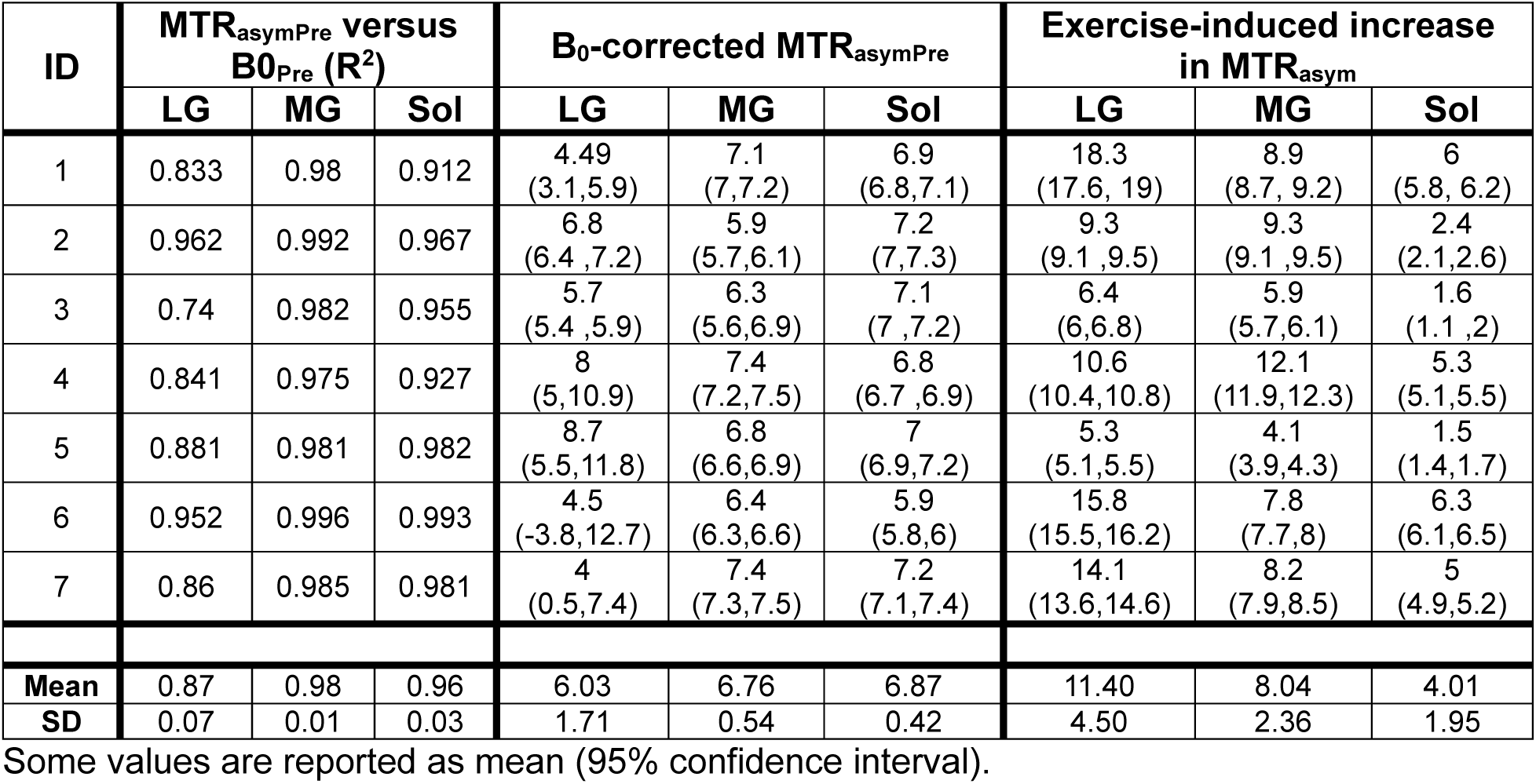
Metrics derived from OXCEST method.

### 4.3 Comparison of OXCEST-derived Cr and 31P-MRS-derived PCr recovery

**Figure 3** shows the comparison between OXCEST-derived post-exercise Cr recovery kinetics and the ground truth post-exercise PCr recovery kinetics in the same subject. Mono-exponential curves were fitted to the corresponding Cr and PCr recovery profiles. The T_PCr_ measured by 31P-MRS was 32s. Using the OXCEST approach, T_Cr_ values were 27s in LG, 25s in MG, and 29s in Sol. Figure 3C illustrates the fitted post-exercise PCr recovery curve alongside the fitted average post-exercise Cr recovery curve from LG and MG, demonstrating strong agreement between OXCEST and 31P-MRS. T_PCr_ and T_Cr_ for all subjects in this study are summarized in **Table 2**. The average T_PCr_ measured was 51 ± 15s. while the T_Cr_ values were 56 ± 19s (LG), 58 ± 29s (MG), and 52 ± 33s (Sol). The combined T_Cr_ of LG and MG showed a significant correlation with T_PCr_ (R^2^ = 0.83, *P* = 0.005) **(Figure 4)**. A CCC value of 0.81 was found between T_Cr_ and T_PCr_, suggesting good agreement.^36^ Representative Sol recovery curves of subjects 5 and 7 with very low T_Cr_ values are in **Figure S4**. Additionally, exercise did not induce any appreciable changes in pH values in six of the seven subjects, while subject 2 exhibited a pH change of 0.7.

**Figure 3.**
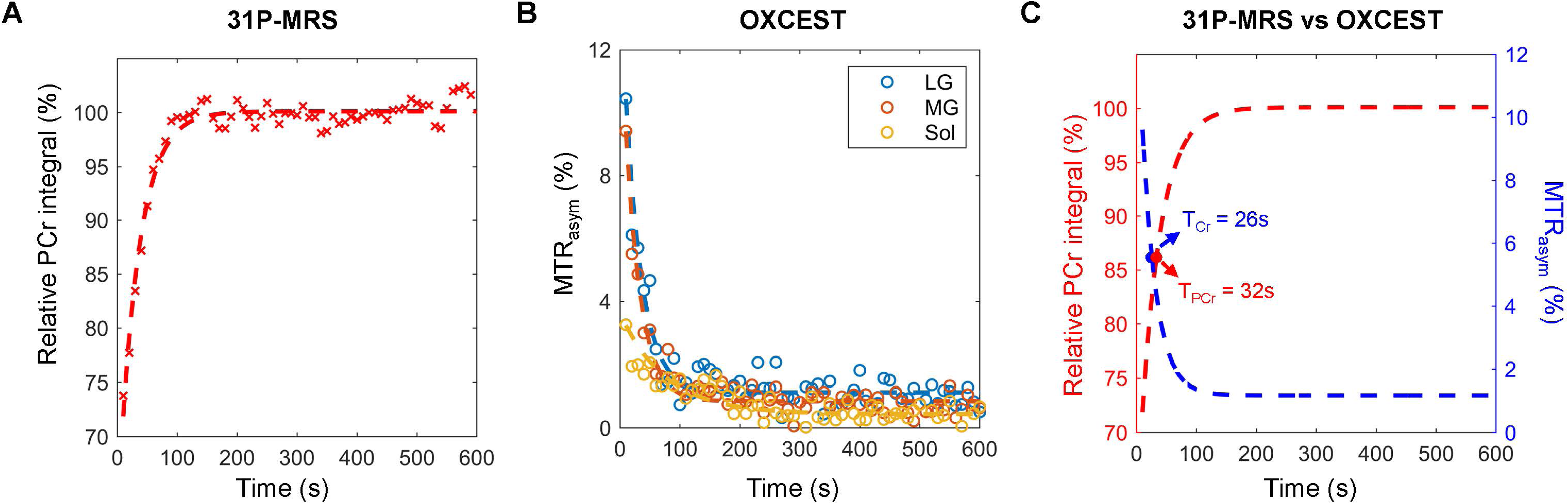
Comparison of post-exercise phosphocreatine (PCr) and creatine (Cr) recovery kinetics measured using 31P-Magnetic Resonance Spectroscopy (31P-MRS) and Oxidative Phosphorylation Chemical Exchange Saturation Transfer (OXCEST), respectively, in a representative subject. (A) Non-localized 31P-MRS-derived PCr signal demonstrates mono-exponential post-exercise growth. (B) OXCEST-derived Cr signal exhibits mono-exponential decay in all three muscles: lateral gastrocnemius (LG), medial gastrocnemius (MG), and soleus (Sol). (C) Mono-exponential curves are fitted to the PCr recovery and to the mean Cr recovery of LG and MG. The PCr recovery time constant derived from 31P-MRS (T_PCr_ = 32s) closely matched the mean Cr recovery time constant (T_Cr_ = 26s) from OXCEST. The faster B_0_ correction in OXCEST enables higher temporal resolution, allowing for detailed capture of early post-exercise dynamics and improved accuracy of kinetic modeling.

**Figure 4.**
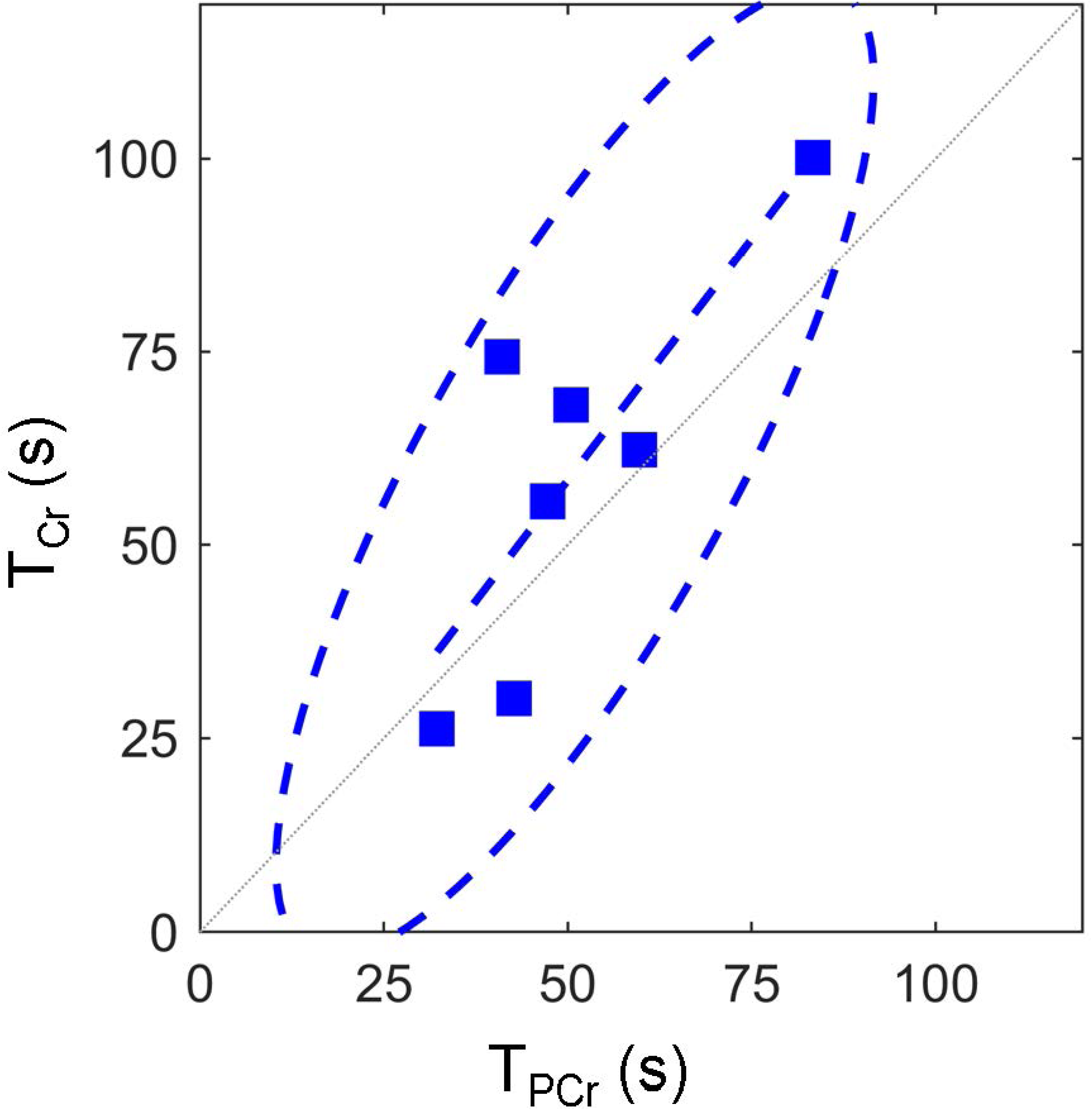
Regression analysis comparing creatine (Cr) recovery time constant (T_Cr_) measured using Oxidative Phosphorylation Chemical Exchange Saturation Transfer (OXCEST) with phosphocreatine (PCr) recovery time constant (T_PCr_) determined using 31P-Magnetic Resonance Spectroscopy (31P-MRS). A significant linear correlation was observed between T_Cr_ and T_PCr_ (R² = 0.83, *P* = 0.005), indicating strong agreement between the two measurements. Dashed ellipse represents the 95% confidence region.

**Table 2:** Recovery time constants derived from OXCEST and 31P-MRS (n = 7).

| ID | 31P-MRS T<sub>PCr</sub>(s) | OXCEST T<sub>Cr</sub>(s) |  |  |
| --- | --- | --- | --- | --- |
|  |  | LG | MG | Sol |
| 1 | 41.06 | 46.15 | 66.54 | 90.4 |
| 2 | 32.24 | 27.19 | 25.27 | 29.33 |
| 3 | 50.43 | 71.04 | 65.23 | 72.46 |
| 4 | 42.74 | 38.94 | 21.37 | 96.52 |
| 5 | 83.28 | 87.96 | 112.39 | 14.56 |
| 6 | 47.24 | 65.95 | 45.34 | 49.84 |
| 7 | 59.77 | 56.66 | 67.92 | 11.75 |
| Mean | 50.97 | 56.27 | 57.72 | 52.12 |
| SD | 15.36 | 19.14 | 28.67 | 32.55 |

## 5 DISCUSSION

In this study, we introduced OXCEST, a novel imaging method for mapping skeletal muscle OXPHOS by applying fast B_0_ correction approach for accurately estimating post-exercise Cr recovery kinetics. OXCEST approach further demonstrated strong agreement with the gold-standard technique, 31P-MRS.

By sampling only two frequency offsets, OXCEST generates B_0_-corrected, Cr-specific MTR_asym_ images, enabling the capture of post-exercise Cr recovery dynamics at very high temporal resolution. This is critical, as literature reports T_PCr_ values ranging from 20-60s in healthy adults,^11,37^ implying that approximately 63% of post-exercise PCr recovery occurs within the first T_PCr_ seconds. Using multiple frequency offsets to compute MTR_asym_ at a single time point, particularly during the post-exercise Cr recovery phase, would inadvertently capture exponentially changing Cr concentrations across offsets, potentially confounding the interpretation of Z-spectra. Given that PCr resynthesis is stoichiometrically coupled with a corresponding decrease in free Cr, capturing the rapid Cr dynamics in muscle demands high temporal resolution. In our study, OXCEST-derived T_Cr_ was significantly correlated with and showed good agreement with 31P-MRS-derived T_PCr_, a marker of mitochondrial OXPHOS capacity. Additionally, the high R² values for the mono-exponential fits in both LG and MG indicate that the model explains over 97% of the variance in the Cr recovery data, supporting the appropriateness of the mono-exponential model for describing post-exercise Cr dynamics. Therefore, OXCEST approach serves as a valuable complement to 31P-MRS technique and offers a non-invasive tool for quantifying skeletal-muscle energetics in clinical settings.

We found that the second-order polynomial relationship between MTR_asymPre_ and ΔB_0Pre_ closely matched previously reported findings in a pH imaging study in rats.^23^ In our study, this fit yielded strong R^2^ values within the observed ΔB_0Pre_ range of −0.25 to +0.25 ppm, indicating robustness across modest B₀ inhomogeneity. We postulate that the proposed B_0_-correction approach in this study will be valid as long as the pre- and post-exercise B_0_ are within moderate drifts (0.25 ppm) away from label frequency of 1.8 ppm. Under these conditions, MTR_asym_ can still be reliably estimated using the proposed theory. However, in the absence of shim, large ΔB_0_ values may lead to DS-affected CEST signal, compromising the validity of the relationship. Although the second-order polynomial fit adequately modeled the relationship between MTR_asymPre_ and B_0Pre_ across all muscles in the three subjects, higher-order fits may need to be explored where a second order model proves insufficient, possibly due to very narrow Cr peak.

We found the net MTR_asymPre_ values for each muscle group to be approximately 6-7%. Reliable estimation of this parameter depends on the presence of some regions within the muscle where ΔB_0_ values are close to zero. If ΔB_0_ values in a muscle are confined to a narrow range far from 0 ppm, the estimate of the y-intercept c_0_ in Equation 2 will have high variability, resulting in wide 95% CIs. We found that MG and Sol consistently met this criterion, having a fraction of their total ROI with ΔB_0_ values close to 0 ppm in all seven subjects. In contrast, the three LG cases that did not meet this criterion were those in which global shimming was performed instead of localized shimming. CrCEST studies in the literature have reported similar MTR_asymPre_ values,^18–20,27,32,38^ supporting the validity of our OXCEST-based quantification. The exercise-induced increase in MTR_asymΔCr_ was found to be highest in LG, followed by MG and Sol, consistent with previous findings.^18,19,27,32,38^ The consistently narrow 95% CIs for these measurements further indicate that OXCEST provides robust MTR_asymPost_ maps and that initial shimming was not a limiting factor. Although the exercise-induced increase in MTR_asymΔCr_ differed markedly among muscles, the mean T_Cr_ remained similar across subjects. This is not contradictory because the two parameters probe separate phases of muscle bioenergetics. ΔMTR_asym_ reflects immediate activation by capturing the extent of phosphocreatine depletion and creatine accumulation during the exercise bout. The magnitudes of the increase are consistent with their relative recruitment during plantar flexion at a given workload. In contrast, T_Cr_ represents post-exercise recovery, governed primarily by mitochondrial capacity to resynthesize PCr (and concomitantly rephosphorylate Cr) independent of the magnitude of the preceding depletion. Thus, ΔMTR_asym_ and T_Cr_ offer complementary information, activation versus recovery, highlighting the complexity of muscle bioenergetics rather than a methodological inconsistency. Substantial inter-subject and inter-muscle variability in both activation and recovery further emphasizes the need to interpret group means with caution. Given its relatively low involvement during the exercise protocol,^32^ Sol in some subjects exhibited T_Cr_ values that differed substantially from those of the LG and MG. Although the AT muscle is generally considered negative control in plantar flexion exercises, both T_Cr_ values and post-exercise increases in MTR_asym_ in our cohort showed high inter-subject variability. While one study reported less than 1% increase in AT MTR_asym_ following mild exercise,^19^ other studies, including unpublished data from our group, have demonstrated measurable AT activation during plantar flexion.^32,39^

The first minute post-exercise is particularly critical for accurately capturing the mono-exponential Cr dynamics. In healthy skeletal muscle, T_PCr_ ranges between 20-60s.^11,37^ Given the exponential nature of PCr recovery, a muscle with T_PCr_ = 20 s recovers ∼95% of PCr by 1 minute post-exercise, whereas a muscle with T_PCr_ = 60 s recovers ∼63% by the same time. This highlights the advantage of OXCEST’s expedient B_0_ correction algorithm, which supports 10s temporal resolution, allowing detailed sampling of the steep early recovery phase. The strong correlation between OXCEST-derived T_Cr_ and 31P-MRS-derived T_PCr_ supports the accuracy of this method.

In one study involving strenuous exercise,^40^ elevated T_2_ relaxation times were inversely correlated with T_PCr_. Since PCr, and Cr are stoichiometrically coupled, any factor affecting PCr kinetics is expected to influence Cr kinetics as well. In the absence of full z-spectra, it is difficult to assess the extent of overlap between T_2_-influenced direct water saturation and the Cr pool at +1.8 ppm. A previous study has shown that T_2_ did not change appreciably after mild exercise.^19^ While participants in our study also performed mild exercise, we did not investigate the potential effects of T_2_* on MTR_asym_.

Acquiring pre-exercise MTR_sym_ values remain important for several reasons, as is also standard practice in conventional CrCEST and 31P-MRS protocols. First, in individuals with neuromuscular disorders who may be unable to complete exercise or remain still during the post-exercise phase, a pre-exercise baseline serves as a critical reference and aids in Cr mapping. Second, lactate accumulation can lead to persistent changes in pH, which may not return to baseline quickly and can further complicate interpretation of recovery dynamics. This is why we recommend using the pre-exercise MTR_asym_ values to estimate the fitting coefficients (c_0_, c_1_, c_2_) in OXCEST.

This study has certain limitations. We used WASSR approach to generate ΔB₀ maps pre- and post-exercise.^21^ A foundational assumption of the OXCEST framework is that the B_0_ inhomogeneity remains stable post-exercise. However, gradual B_0_ field drift may still occur due to heating of scanner components during gradient-intensive MR sequences, a phenomenon observed in NOE experiments.^24,41^ Similar local magnetic field drifts are a well-recognized issue in functional MRI studies.^42^ Additionally, post-exercise changes such as muscle recovery or subtle calf movement can affect the B_0_, introducing errors in correction. To mitigate such effects, an interleaved gradient echo-based B_0_ mapping sequence could be implemented alongside the CEST acquisition to regress out the effect of such frequency drifts.^43^ Acquiring dynamic ΔB_0Post_ for each frequency offset pairs would enable accurate computation of MTR_asymΔB0_ (Equation 3), and subsequently MTR_asymΔCr_ and T_Cr_, making OXCEST more robust to motion and the resulting B_0_ fluctuations over time. Secondly, our manuscript included data from healthy volunteers only. In subjects with neuromuscular diseases, the lactate accumulation can significantly change the pH, which will change the exchange rates, resulting in underestimation of the CEST contrast.^20^ Notably, pH-specific changes have been accounted for in a recent report by Philip Zhe Sun through the inclusion of an additive correction term.^23^ Finally, OXCEST and 31P-MRS recovery data were acquired during two separate sessions conducted on different days. Although care was taken to ensure that exercise bouts were consistent across sessions, inter-session variability may still influence direct comparisons. A large-scale study with simultaneous monitoring of PCr and Cr recovery kinetics is needed to confirm the reproducibility and robustness of these findings.

## 6 CONCLUSION

In this study, we introduced OXCEST, an advanced technique for mapping skeletal muscle OXPHOS by quantifying CEST effects of Cr using acquisitions at two frequency offsets. We demonstrated its utility by mapping dynamic changes in skeletal muscle metabolism post-exercise in healthy participants and comparing the results with the gold standard 31P-MRS technique. This validation highlights OXCEST’s potential for clinical translation as a potential non-invasive alternative for assessing in vivo muscle mitochondrial OXPHOS. Furthermore, the underlying principles of OXCEST are broadly applicable and adapted to other CEST-based imaging applications, particularly those requiring high temporal resolution following a physiological or pharmacological intervention.

## ACKNOWLEDGEMENTS

The authors thank Ms. Jennifer Valley and Dr. Mitchell Taylor for their technical support and Ms. Pamela Adede for her assistance with participant recruitment.

## DATA AVAILABILITY STATEMENT

Data will be made available upon request.

