## Supplemental Data for "A novel CEST-based approach for reliably assessing skeletal muscle oxidative phosphorylation: OXCEST"

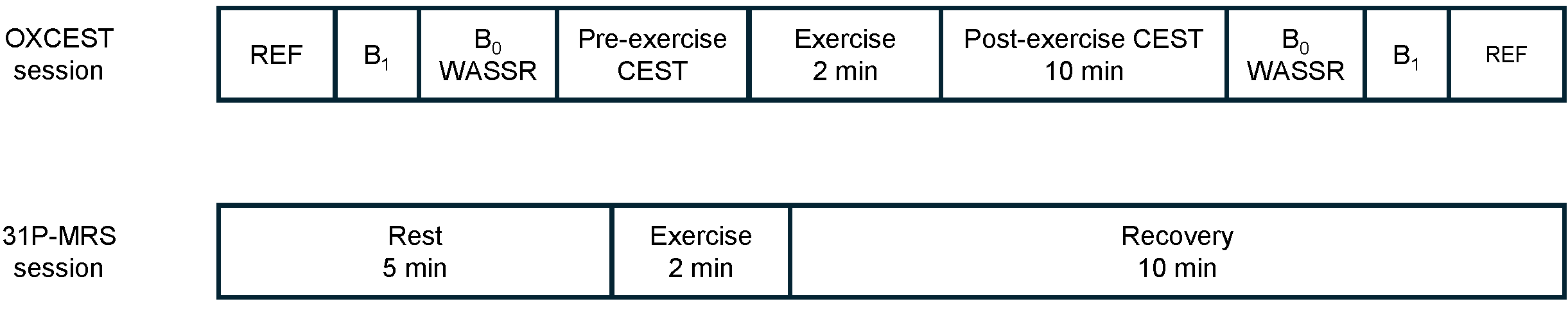

**Figure S1.** Schematic of the imaging protocol for Oxidative Phosphorylation Chemical Exchange Saturation Transfer (OXCEST) and 31P-Magnetic Resonance Spectroscopy (31P-MRS) for each subject. MRI scans were performed over two separate sessions on different days. During each session, participants performed mild plantar flexion exercise inside the scanner for 2 minutes at a rate of 45 flexions per minute.

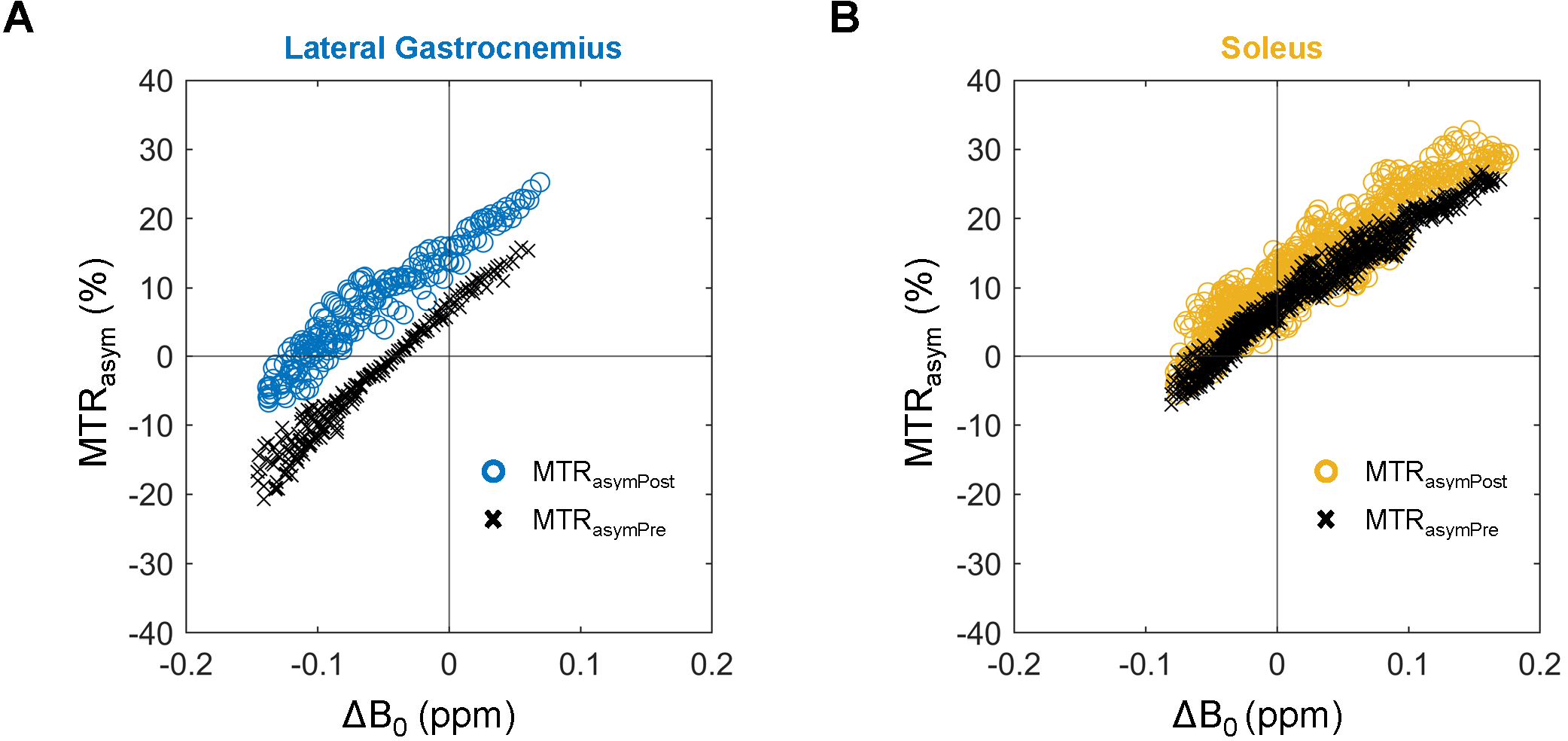

**Figure S2:** Pixel-wise scatterplot of Oxidative Phosphorylation Chemical Exchange Saturation Transfer (OXCEST) magnetization transfer ratio asymmetry (MTR_asym_) versus main magnetic field inhomogeneity (ΔB_0_) in the lateral gastrocnemius (panel A) and soleus (panel B) of the same representative subject shown in Figure 1. Black crosses indicate pre-exercise MTR_asym_ (MTR_asymPre_); colored hollow circles represent post-exercise MTR_asym_ (MTR_asymPost_).

| **(A)** | 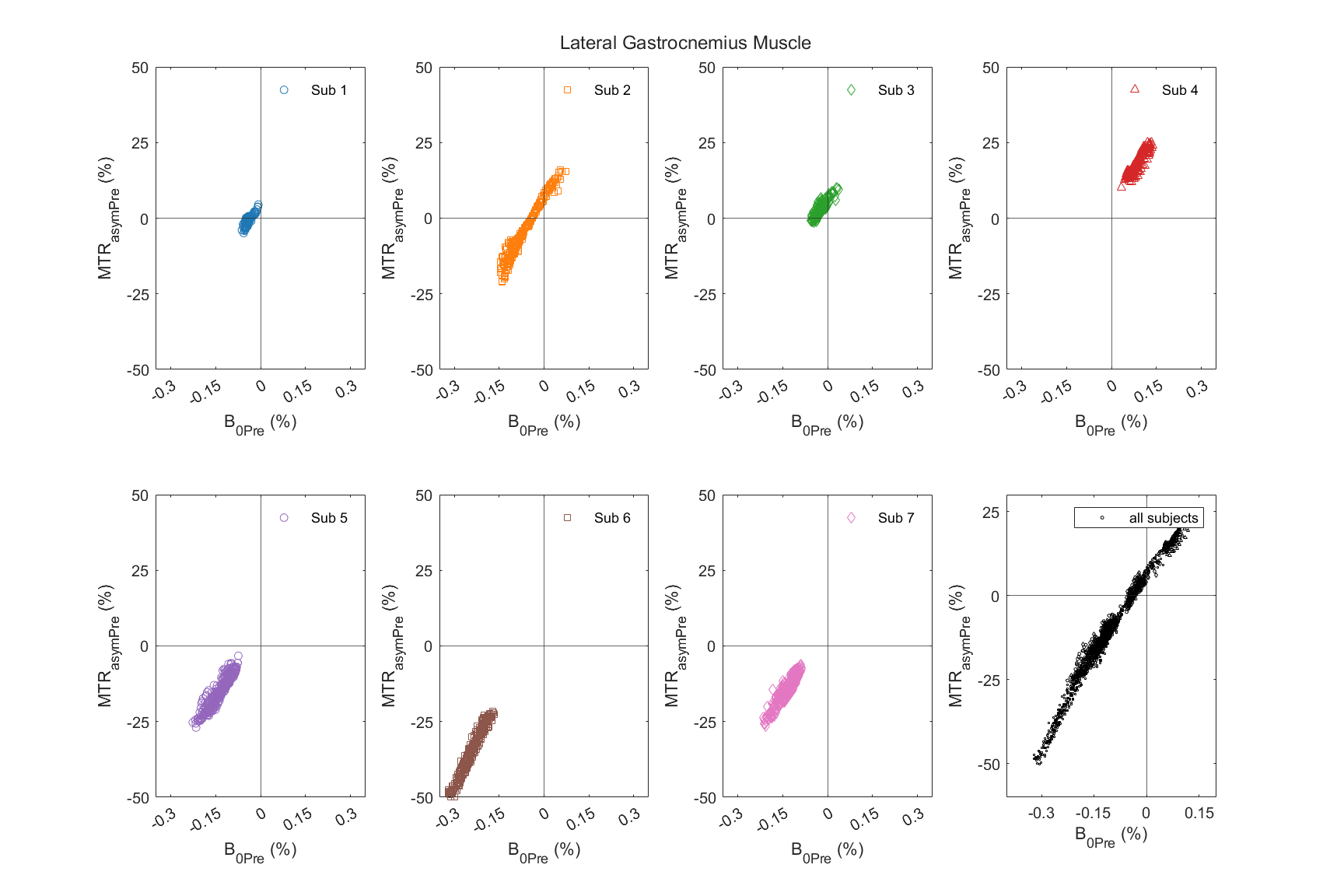 |
| --- | --- |
| **(B)** | 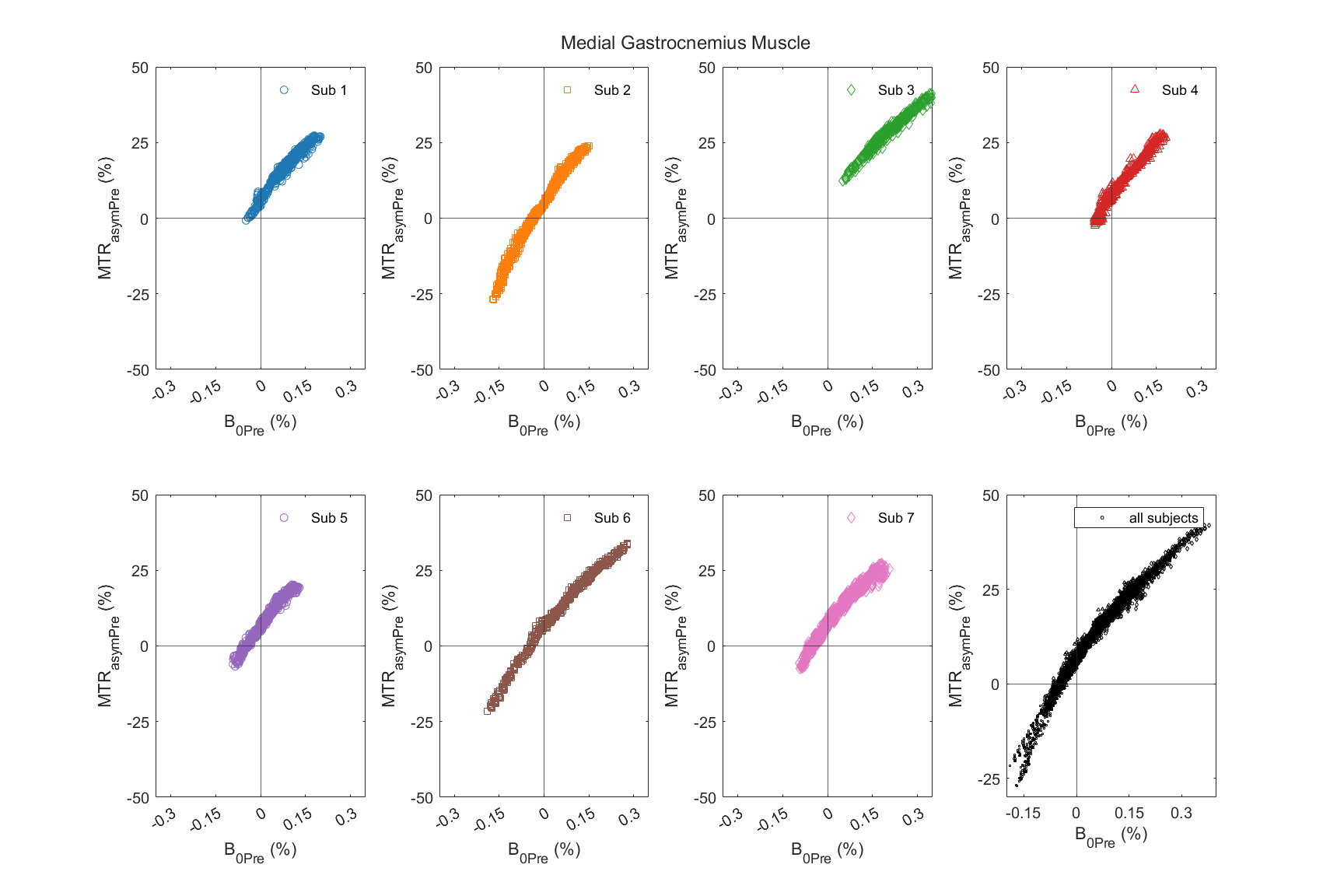 |
| **(C)** | 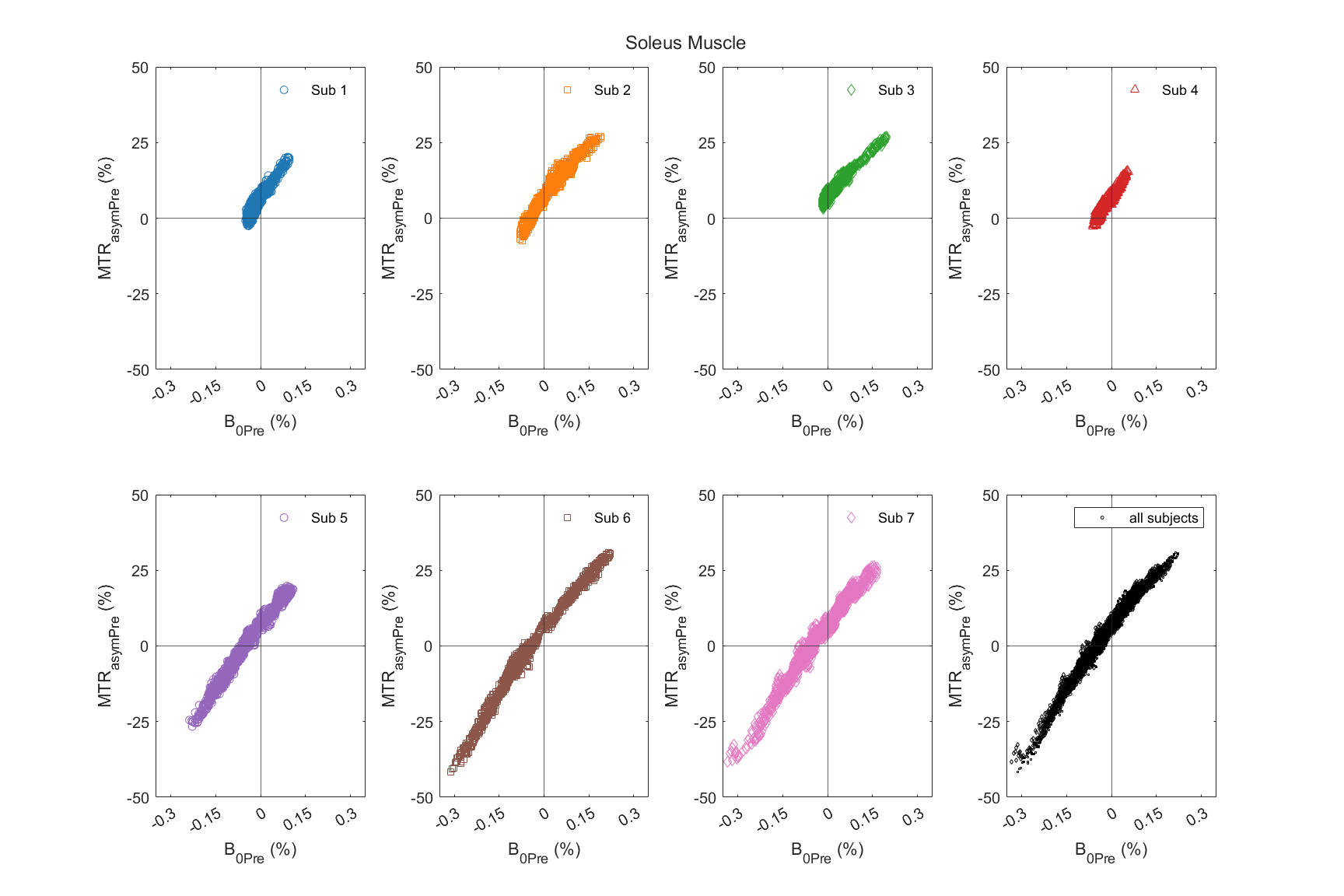 |
| **Figure S3.** Pre-exercise magnetic transfer ratio asymmetry (MTR_asymPre_) modelled against pre-exercise B_0_ inhomogeneity (ΔB_0Pre_) using 2^nd^ order polynomial in Oxidative Phosphorylation Chemical Exchange Saturation Transfer (OXCEST). | |

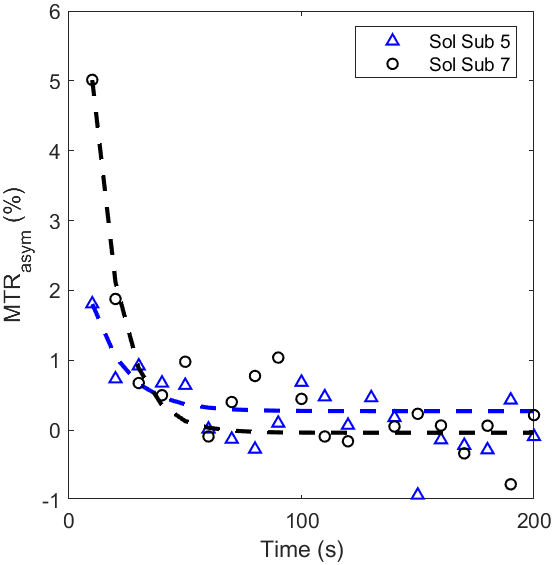

**Figure S4.** Representative fit curves for two subjects with unusually short creatine recovery time (T_Cr_) of less than 15s.

**Table S1:** Fitted coefficients of OXCEST that modeled MTR_asymPre_ against B_0Pre_ in lateral gastrocnemius (LG), medial gastrocnemius (MG), and soleus (Sol) muscles

| **ID** | **LG** | | | **MG** | | | **Sol** | | |
| --- | --- | --- | --- | --- | --- | --- | --- | --- | --- |
|  | **c_2_** | **c_1_** | **c_0_** | **c_2_** | **c_1_** | **c_0_** | **c_2_** | **c_1_** | **c_0_** |
| 1 | 54.77 | 136.9 | 4.49 | -202.9 | 145.8 | 7.094 | -184.9 | 150.7 | 6.934 |
| 2 | -135 | 153.4 | 6.764 | -191.3 | 152.3 | 5.91 | -222.9 | 144.1 | 7.153 |
| 3 | -120.3 | 107.7 | 5.683 | -86.73 | 127.8 | 6.287 | -105.5 | 119.4 | 7.072 |
| 4 | 240.7 | 93.96 | 7.977 | -138.2 | 137.5 | 7.353 | 162.5 | 139.4 | 6.815 |
| 5 | 251 | 209.4 | 8.692 | -180.3 | 133.6 | 6.763 | -78.76 | 130.3 | 7.03 |
| 6 | -146.6 | 126.2 | 4.481 | -118 | 128.1 | 6.432 | -89.52 | 128.9 | 5.908 |
| 7 | -40.64 | 128.3 | 3.959 | -218.4 | 136.5 | 7.384 | -68.4 | 127.9 | 7.231 |

**Table S2:** Distribution of B_0_ coverage (in ppm) during pre- and post-exercise phase in lateral gastrocnemius (LG), medial gastrocnemius (MG), and soleus (Sol) muscles

| **ID** | **LG** | | **MG** | | **Sol** | |
| --- | --- | --- | --- | --- | --- | --- |
|  | **Pre-exercise** | **Post-exercise** | **Pre-exercise** | **Post-exercise** | **Pre-exercise** | **Post-exercise** |
| 1 | (-0.06, 0) | (-0.09, -0.04) | (-0.05, 0.2) | (-0.08, 0.2) | (-0.05, 0.1) | (-0.07, 0.07) |
| 2 | (-0.15, 0.07) | (-0.14, 0.08) | (-0.17, 0.15) | (-0.18, 0.15) | (-0.08, 0.19) | (-0.08, 0.21) |
| 3 | (-0.06, 0.04) | (-0.08, 0.03) | (0.05, 0.38) | (0.02, 0.35) | (-0.02, 0.2) | (-0.02, 0.2) |
| 4 | (0.03, 0.14) | (-0.01, 0.1) | (-0.06, 0.18) | (-0.09, 0.16) | (-0.06, 0.06) | (-0.09, 0.03) |
| 5 | (-0.23, -0.07) | (-0.23, -0.09) | (-0.09, 0.13) | (-0.12, 0.12) | (-0.24, 0.11) | (-0.25, 0.11) |
| 6 | (-0.32, -0.17) | (-0.33, -0.18) | (-0.19, 0.28) | (-0.22, 0.26) | (-0.31, 0.22) | (-0.35, 0.24) |
| 7 | (-0.21, -0.09) | (-0.24, -0.06) | (-0.1, 0.2) | (-0.12, 0.225) | (-0.33, 0.17) | (-0.33, 0.18) |

All data reported as (min, max)

**Table S3:** Noise level and goodness-of-fit metrics for mono-exponential Creatine recovery in lateral gastrocnemius (LG), medial gastrocnemius (MG), and soleus (Sol) muscles

| **ID** | **Noise SD (% of exercise-induced MTR_asym_ increase)** | | | **RMSE (R^2^) of mono-exponential fit** | | |
| --- | --- | --- | --- | --- | --- | --- |
|  | **LG** | **MG** | **Sol** | **LG** | **MG** | **Sol** |
| 1 | 6.6 | 6.9 | 7.1 | 1.23 (0.988) | 0.64 (0.988) | 0.53 (0.981) |
| 2 | 5.4 | 3.9 | 18.4 | 0.51 (0.995) | 0.37 (0.997) | 0.38 (0.968) |
| 3 | 15.6 | 7.4 | 27.9 | 0.73 (0.977) | 0.39 (0.992) | 0.41 (0.816) |
| 4 | 5.0 | 4.6 | 7.7 | 0.54 (0.996) | 0.56 (0.997) | 0.42 (0.988) |
| 5 | 9.4 | 11.5 | 30.1 | 0.5 (0.983) | 0.48 (0.973) | 0.47 (0.869) |
| 6 | 5.7 | 5.3 | 9.4 | 0.92 (0.994) | 0.42 (0.995) | 0.61 (0.984) |
| 7 | 9.4 | 9.4 | 9.7 | 1.3 (0.984) | 0.79 (0.983) | 0.5 (0.985) |

SD: standard deviation; RMSE: root mean squared error; R^2^: coefficient of determination

**Table S4:** Effect of B_0_ correction on MTR_asym_ in lateral gastrocnemius (LG), medial gastrocnemius (MG), and soleus (Sol) muscles

| **ID** | **MTR_asymPre_** | | | | | | **Exercise-induced increase in MTR_asym_** | | | | | |
| --- | --- | --- | --- | --- | --- | --- | --- | --- | --- | --- | --- | --- |
|  | **Without B_0_-correction** | | | **With B_0_-correction** | | | **Without B_0_-correction** | | | **With B_0_-correction** | | |
|  | **LG** | **MG** | **Sol** | **LG** | **MG** | **Sol** | **LG** | **MG** | **Sol** | **LG** | **MG** | **Sol** |
| 1 | -14.5 | 9.9 | 0.6 | 4.49 | 7.1 | 6.9 | 9.1 | 8 | 4.4 | 18.3 | 8.9 | 6 |
| 2 | -20.3 | -2 | -0.64 | 6.8 | 5.9 | 7.2 | 10.3 | 8.2 | 2.9 | 9.3 | 9.3 | 2.4 |
| 3 | 3.8 | 28.7 | 12.1 | 5.7 | 6.3 | 7.1 | 5.3 | 6.2 | 0.7 | 6.4 | 5.9 | 1.6 |
| 4 | 17.9 | 12.9 | 5.9 | 8 | 7.4 | 6.8 | 7.2 | 9.3 | 2.4 | 10.6 | 12.1 | 5.3 |
| 5 | -14 | 8.9 | -0.2 | 8.7 | 6.8 | 7 | 3.1 | 2.7 | 1.7 | 5.3 | 4.1 | 1.5 |
| 6 | -34 | 9.9 | -2.4 | 4.5 | 6.4 | 5.9 | 14.6 | 5.3 | 6.8 | 15.8 | 7.8 | 6.3 |
| 7 | -13.3 | 13.1 | 1 | 4 | 7.4 | 7.2 | 12.4 | 4.1 | 4.8 | 14.1 | 8.2 | 5 |

**Table S5.** OXCEST metrics in anterior tibialis muscle across all subjects (n=7)

| **ID** | **T_Cr_ (s)** | **Exercise-induced increase in MTR_asym_** |
| --- | --- | --- |
| 1 | 93.6 | 5.1 |
| 2 | 49.2 | 1.59 |
| 3 | N/A | 0 |
| 4 | 40.6 | 7.96 |
| 5 | 18.4 | 1.5 |
| 6 | 48.5 | 11.5 |
| 7 | 15.4 | 12.5 |
